# Development of a Whole-Cell Biosensor for β-Lactamase Inhibitor Discovery

**DOI:** 10.1101/2023.07.24.550393

**Authors:** Mitchell A. Jeffs, Rachel A.V. Gray, Prameet M. Sheth, Christopher T. Lohans

## Abstract

The clinical utility of the β-lactam antibiotics has been endangered by the production of β-lactamases by β-lactam-resistant pathogenic bacteria such as *Escherichia coli, Pseudomonas aeruginosa* and *Acinetobacter baumannii*. Collectively, these enzymes can degrade every clinically available β-lactam, jeopardizing antimicrobial therapy. Although extensive efforts have been made to develop β-lactamase inhibitors, inhibitor-resistant β-lactamases emerge rapidly. In addition, there are currently no clinically available inhibitors against the metallo-β-lactamases, a group of β-lactamases of great global concern. To further inhibitor discovery efforts, new assays are required to assess inhibitor efficacy, particularly in a cellular context. We report the development of a whole-cell *E. coli* biosensor which can quantify β-lactamase inhibition in a cellular context. Upon administration of an effective inhibitor, a β-lactam is rescued from β- lactamase-catalyzed degradation, resulting in the emission of a luminescent signal by the biosensor. This platform was validated using a panel of clinically relevant β-lactamases and was applied to quantitatively study the potency of a selection of currently used and reported β-lactamase inhibitors. This rapid method can account for factors like membrane permeability and can be employed to identify new β-lactamase inhibitors.

## INTRODUCTION

Currently, β-lactams are the most frequently prescribed class of antibiotics worldwide [1]. However, bacterial pathogens have acquired resistance mechanisms to overcome these medications, rendering them ineffective. Among Gram-negative bacteria, β-lactamases are one of the most prevalent causes of resistance [2]. These enzymes degrade β-lactams by hydrolyzing the β-lactam ring, a feature required for antibacterial activity (Fig. 1A). β-lactamases are classified according to their mechanism: serine β-lactamases (SBLs) employ a catalytic serine residue, while metallo-β-lactamases (MBLs) activate a hydrolytic water molecule using one or more zinc ions [3]. SBLs and MBLs are produced by bacterial pathogens of significance to human health, including *Pseudomonas aeruginosa*, *Acinetobacter baumannii*, *Klebsiella pneumoniae*, among others [4].

**Fig. 1.**
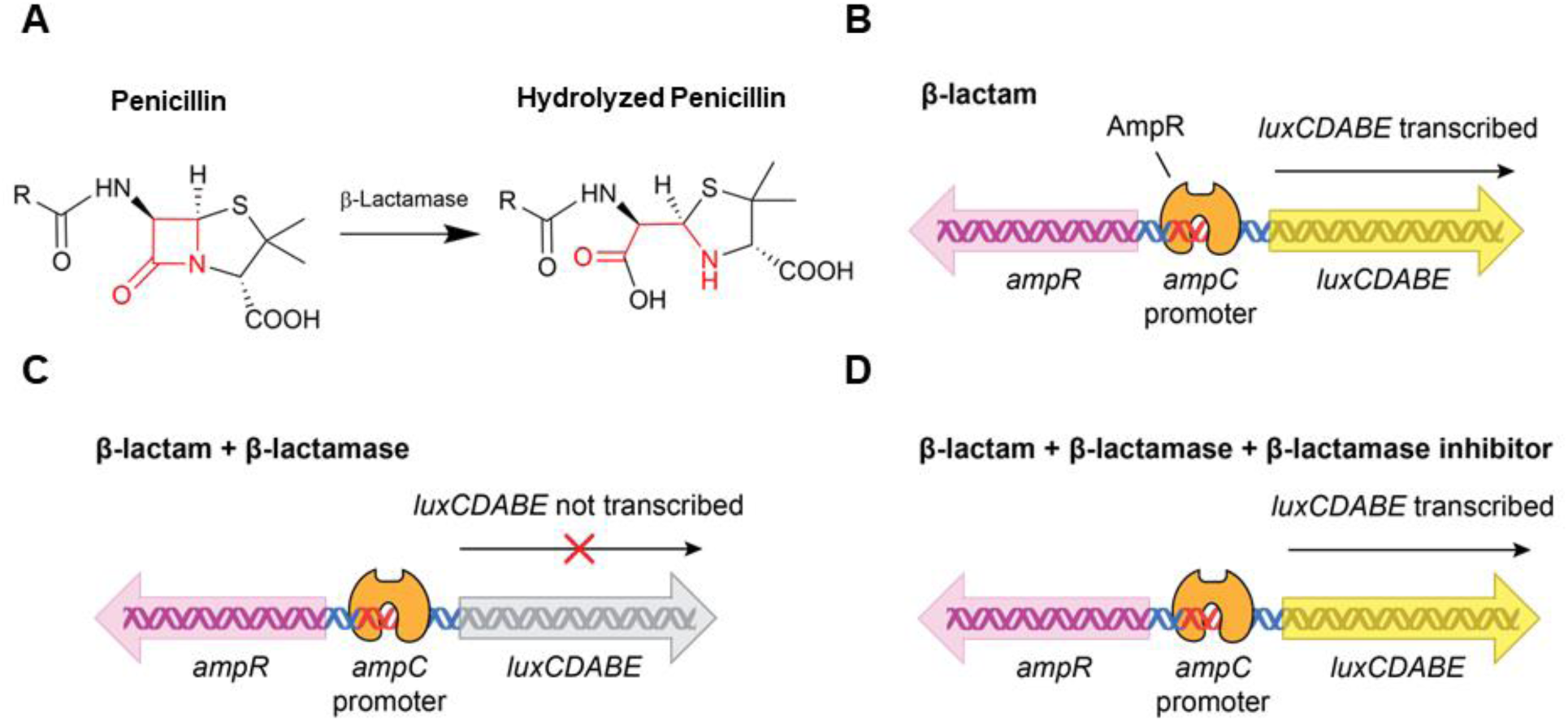
Rationale for the developed biosensor. (A) General reaction scheme depicting β-lactam hydrolysis by a β-lactamase. (B) Exposure of *E. coli* cells transformed with pAMPLUX to a β- lactam induces *lux* operon transcription, resulting in luminescence. (C) If the β-lactam is degraded by a β-lactamase, the amount of luminescence is reduced. (D) Upon administration of an effective β-lactamase inhibitor, luminescence is restored.

Despite successful efforts to develop β-lactamase inhibitors that can be administered with β-lactams (e.g., the SBL inhibitors clavulanic acid, tazobactam, and avibactam), use of these inhibitors was soon followed by the identification of β-lactamases that were less susceptible to them [5–7]. For example, a β-lactamase variant that was resistant to avibactam was reported in 2015, the same year this inhibitor was approved for clinical use [6]. To make matters worse, there are currently no MBL inhibitors available for therapeutic purposes [8]. Although there are promising MBL inhibitors in clinical testing (e.g., taniborbactam), resistant MBL variants have already been detected [9]. Thus, there is a dire need for new β-lactamase inhibitors.

Current methods for identifying β-lactamase inhibitors often employ purified enzymes, monitoring the inhibition of β-lactamase-catalyzed hydrolysis of chromogenic and fluorogenic substrates [10, 11]. However, identifying inhibitors that are active against cellular β-lactamases is vital, as inhibitor potency against purified enzymes often does not reflect potency against cells [12, 13]. Traditionally, growth-based minimum inhibitory concentration (MIC) tests are used to assess inhibitor efficacy against β-lactamase-producing cells, but turbidity measurements lack sensitivity and specificity. To further β-lactamase inhibitor discovery efforts, new screening assays are required. We report the development and validation of a whole-cell biosensor assay that can be applied to the discovery of β-lactamase inhibitors. This approach does not require the use of expensive substrates and is selective for inhibitors that are effective against bacterial cells.

## RESULTS AND DISCUSSION

Our biosensor platform employs the AmpR/AmpC regulatory system from P. aeruginosa [14]. Exposure of *P. aeruginosa* to certain β-lactams elevates production of 1,6- anhydromuropeptides during peptidoglycan catabolism; some of these catabolites bind to AmpR, inducing the expression of the *ampC* β-lactamase gene [15]. We prepared the plasmid pAMPLUX in which the luxCDABE operon is controlled by the *ampC* promoter, such that exposure to a β- lactam antibiotic leads to a luminescent signal (Fig. 1B). Related reporter systems have been applied to study β-lactam biosynthesis and peptidoglycan recycling [16–18]. If the β-lactam antibiotic is degraded by β-lactamase-producing bacteria prior to the addition of biosensor cells, no luminescence would be observed (Fig. 1C). Finally, addition of an effective β-lactamase inhibitor protects the β-lactam, restoring luminescence (Fig. 1D).

The pAMPLUX plasmid was prepared by cloning the lux operon, *ampR*, and the *ampC* promoter into a kan^R^ derivative of pUC19 (Fig. S1). Treatment of *E. coli* BW25113 cells transformed with pAMPLUX with imipenem and amoxicillin (β-lactams which induce the AmpR system) [19, 20] resulted in a strong luminescent signal (Fig. 2A) which was not observed for untransformed cells (Fig. S2). Luminescence increased with antibiotic concentration until the amount of antibiotic was sufficient to kill the biosensor cells (Fig. 2B). Optimization experiments were conducted to determine the optimal incubation temperature and culture density for subsequent assays (Fig. S3, S4). We next tested the ability of the *E. coli* biosensor to monitor β- lactam degradation by β-lactamase-producing cells, beginning with the SBL TEM-116. This enzyme is a penicillinase which hydrolyzes penicillins [e.g., amoxicillin (AMX)]. After pre-treating TEM-116- producing *E. coli* cells with AMX, biosensor cells were added. After two hours of incubation, only low levels of luminescence were observed, indicating AMX degradation by TEM-116 (Fig. 3A). Preincubation of TEM-116 producing *E. coli* with AMX and SBL inhibitors tazobactam (TZB), clavulanic acid (CLAV) or avibactam (AVI) [21–23] prior to the addition of biosensor cells resulted in an increase in luminescence, demonstrating the rescue of AMX (Fig. 3A).

**Fig. 2.**
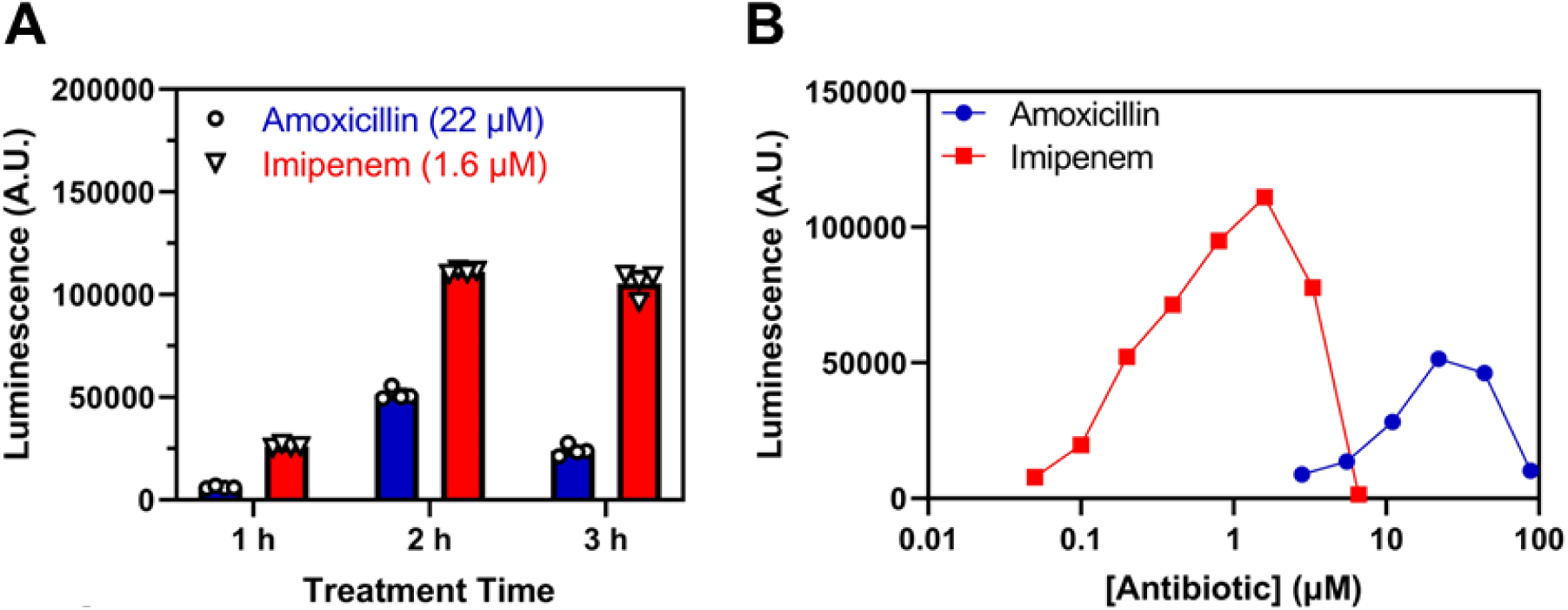
Initial characterization of the biosensor. (A) Treatment of biosensor cells with amoxicillin (AMX) or imipenem (IMI) leads to a time-dependent luminescent signal. (B) Dose response curves for AMX and IMI with biosensor cells, measured two hours post-antibiotic exposure. n=4, error bars indicate S.D.

**Fig. 3.**
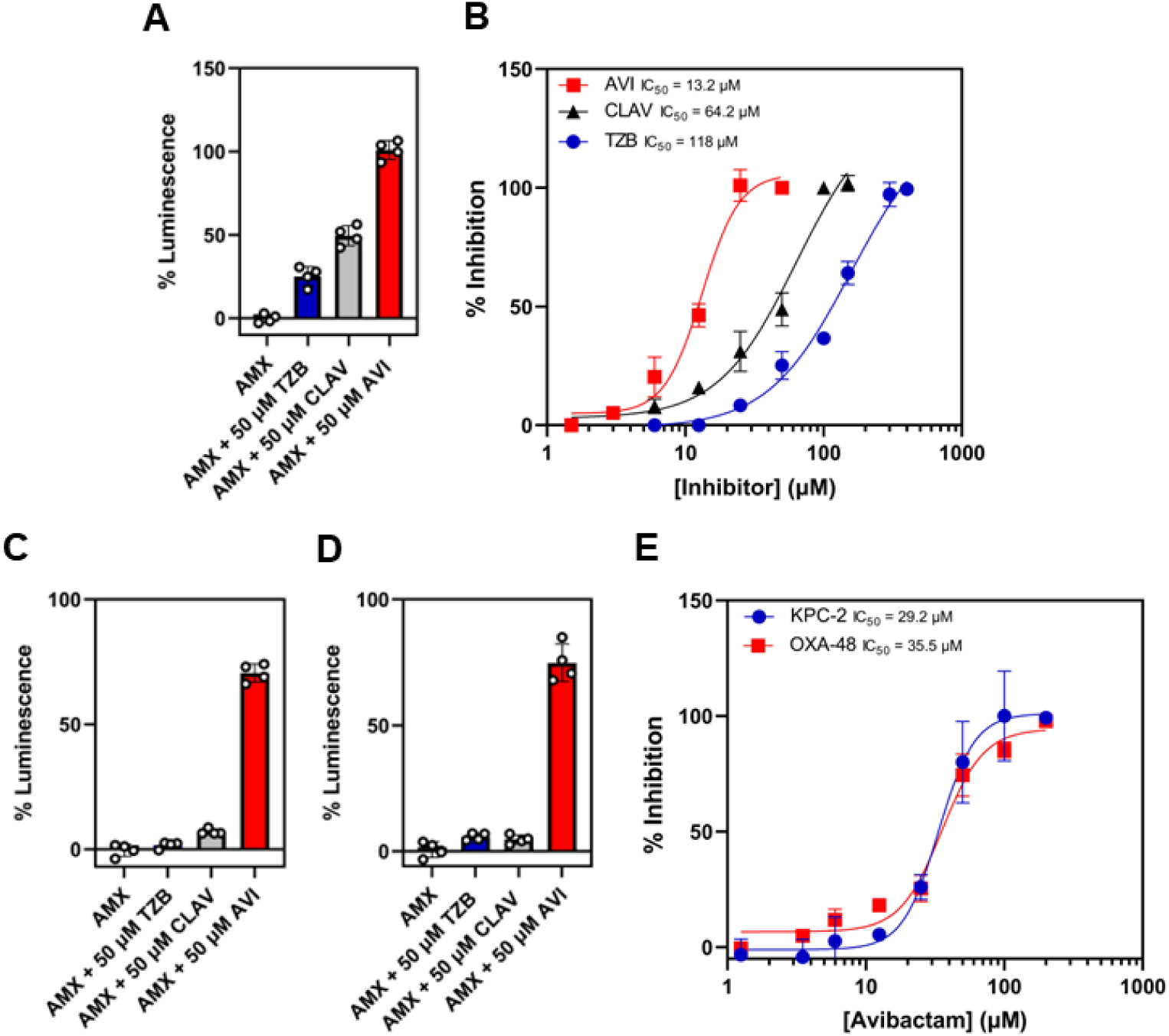
SBL inhibition assays. (A) Single-point TEM-116 inhibition assays and (B) TEM-116 dose response analysis. TEM-116-producing *E. coli* were treated with amoxicillin (AMX, 20 µM) in combination with the indicated concentrations of avibactam (AVI), clavulanic acid (CLAV) or tazobactam (TZB). Single-point inhibition assays for (C) KPC-2 and (D) OXA-48. (E) KPC-2 and OXA-48 dose-response inhibition by avibactam. The same conditions were used as for the TEM- 116 inhibition assays. Luminescence readings were normalized to the sample with the greatest luminescence to determine % inhibition. n=4, error bars indicate S.D.

To expand, we evaluated the relationship between inhibitor concentration and biosensor induction in the presence of a fixed antibiotic concentration (Fig. 3B). As the concentration of inhibitor was increased, the amount of luminescence also increased, indicating less antibiotic degradation by TEM-116. The resulting dose response curves were used to determine effective IC_50_ values for the inhibitors tested (Table S3), quantifying their potencies in a cellular context. These data show that AVI is the most potent inhibitor of TEM-116, consistent with published reports for purified TEM-1 [24]. We verified these results with traditional turbidity-based assays by conducting growth-based experiments under the same conditions as the biosensor assays (Table S3). It should be noted that TZB, AVI, and CLAV induce anhydromuropeptide production and can activate the biosensor in the absence of antibiotic. However, the levels of induction were relatively low at the concentrations tested when compared to their antibiotic partners (Fig. S5) and were accounted for during inhibition studies. The potential inhibition of penicillin-binding proteins (PBPs) by new β-lactamase inhibitors can be assessed by testing the inhibitors against biosensor cells in the absence of β-lactams and β-lactamase-producing cells.

Following the experiments with TEM-116, we extended our inhibition studies to include KPC-2 and OXA-48, SBLs of great clinical significance due in part to their ability to degrade carbapenem antibiotics [25]. While treatment of *E. coli* cells that produce KPC-2 or OXA-48 with AVI rescued AMX, leading to a strong luminescent signal, TZB and CLAV did not inhibit these enzymes under the conditions tested (Fig. 3C, 3D). Dose response analyses indicated that avibactam inhibited KPC-2 and OXA-48 with similar potencies (Fig. 3E) These observations are consistent with previous studies testing these inhibitors against purified enzymes, and also against cells that produce these β-lactamases (Table S3) [26–28].

Next, we expanded our inhibition studies to the MBLs NDM-1, IMP-1, and VIM-2. As there are not currently any MBL inhibitors approved for clinical use, we tested five inhibitors described in the literature: ethylenediaminetetraacetic acid (EDTA), dipicolinic acid (DPA), nitrilotriacetic acid (NTA), captopril (CAP), and embelin (EMB) [29–31]. Our assay results suggest that EDTA is the most potent inhibitor against NDM-1 (Fig. 4A, 4B), followed by EMB, DPA, NTA, and finally CAP. These trends are consistent with published IC_50_ values for these inhibitors against purified NDM-1 [29–32]. While EDTA was also the most effective inhibitor against IMP-1 and VIM-2 (Fig. S6, S7), EMB and NTA exhibited poor activity against IMP-1 in agreement with literature reports [30, 31].

**Fig. 4.**
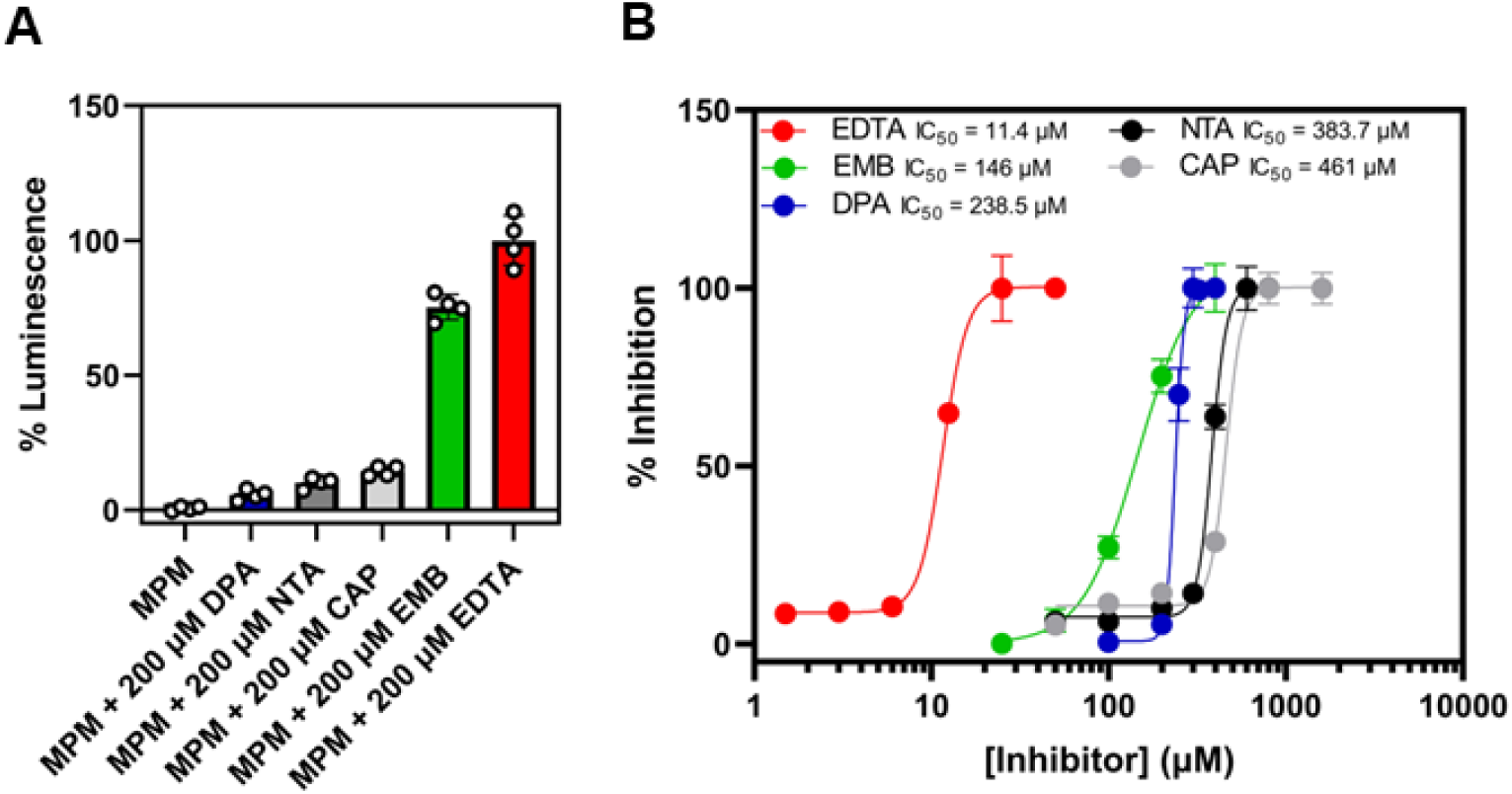
NDM-1 inhibition assays. (A) Single point NDM-1 inhibition and (B) NDM-1 dose response analysis. NDM-1-producing E. coli were treated with meropenem (MPM, 2.5 µM) in combination with the indicated concentrations of ethylenediaminetetraacetic acid (EDTA), dipicolinic acid (DPA), nitrilotriacetic acid (NTA), captopril (CAP) and embelin (EMB). Luminescence readings were normalized to the sample with the greatest luminescence to determine % inhibition. n=4, error bars indicate S.D.

It should be noted that the cellular IC_50_ values obtained in this study may be impacted by several factors. Firstly, the inoculum effect (i.e., an increase in the MIC of an antimicrobial agent when the inoculum size of test bacteria is increased) should be considered [33, 34]. We found that the inoculum of β-lactamase-producing cells can be adjusted (Fig. S8) to identify inhibitors of higher or lower potency. Cellular IC_50_ values could also be impacted by the destabilization of the outer membrane by metal chelating inhibitors (e.g., EDTA), which allows easier passage of small molecules across the cell membrane [35]. This could increase the entry of β-lactams and β- lactamase inhibitors, resulting in increased potency of these agents. β-lactamase inhibitors that also permeabilize the outer membrane (e.g., EDTA) can be identified by assessing luminescence production following treatment of biosensor cells with inhibitor alone, or in combination with β- lactams in the absence of a β-lactamase (Fig. S9).

Beyond the production of β-lactamases, Gram-negative pathogens employ other resistance mechanisms that impact the efficacy of β-lactams and β-lactamase inhibitors (e.g., reduced porin expression, efflux pumps) [36]. While purified enzyme assays cannot account for the presence of other resistance mechanisms, our biosensor platform can evaluate the impact of cellular factors such as decreased permeability. The *E. coli* porin OmpF is used by certain β-lactams for cellular entry, and decreased production of this porin is associated with resistance to β-lactams and β- lactam-β-lactamase inhibitor combinations (e.g., ceftazidime-avibactam) [37, 38] As proof of concept, we used the biosensor to evaluate the efficacy of β-lactamase inhibitors against an E. coli Δ*ompF* strain that produces TEM-116. A notable increase in IC_50_ was observed for both AVI and TZB when administered with AMX against this strain (Fig. 5), highlighting the importance of membrane permeability on the efficacy of β-lactam-β-lactamase inhibitor combinations. Thus, the biosensor could be used to prioritize the identification of β-lactamase inhibitors that function against β-lactamase-producing bacteria that possess additional resistance mechanisms.

**Fig. 5.**
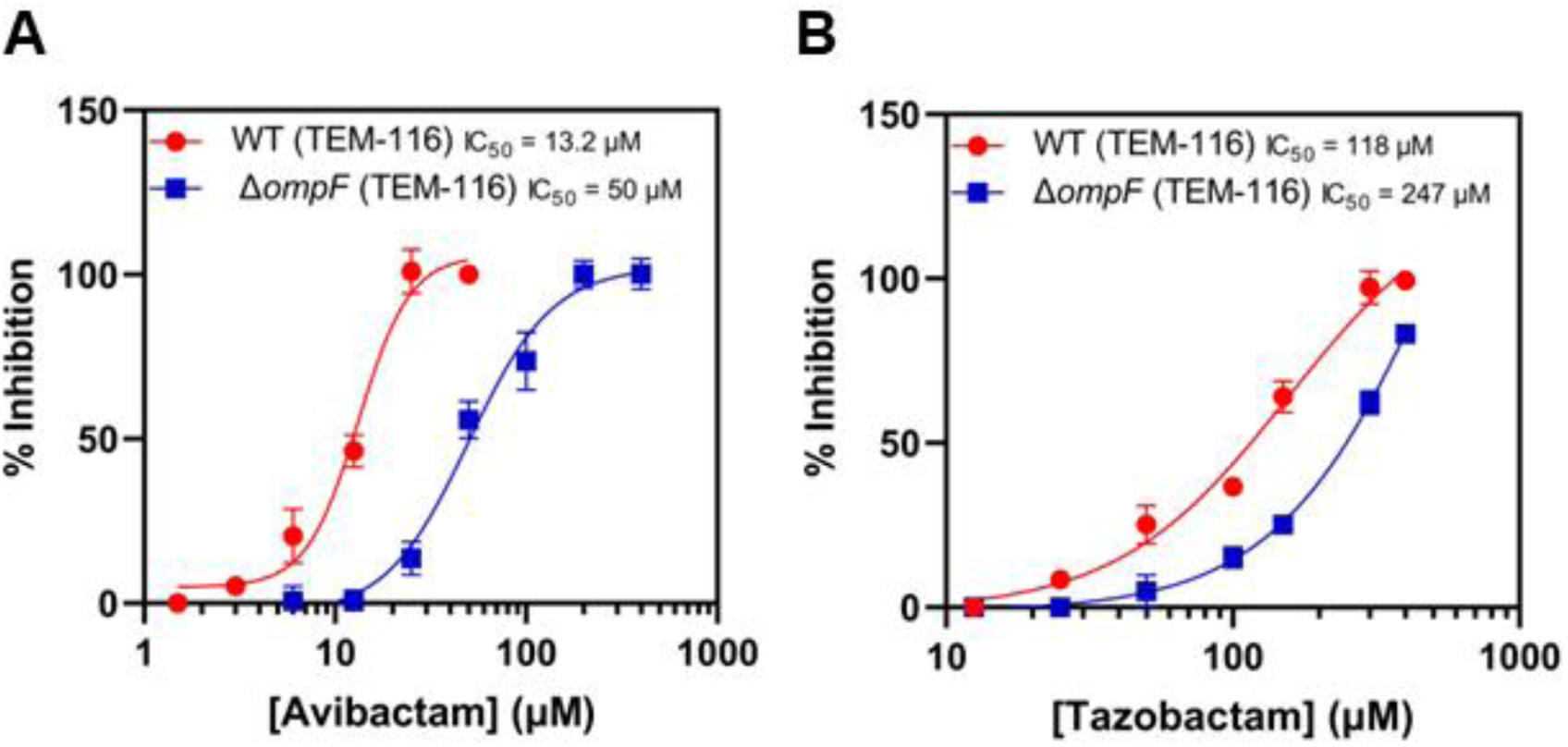
Effects of *E. coli ompF* knockout on TEM-116 inhibition. Dose response analysis of (A) avibactam and (B) tazobactam in combination with amoxicillin (20 µM). Previously determined dose-response curves for the wild-type (WT) strain are shown for comparison. Luminescence readings were normalized to the sample with the greatest luminescence to determine % inhibition. n=4, error bars indicate S.D.

Our results validate the application of this *E. coli* biosensor to the measurement of β- lactamase activity and inhibition in whole cells. Rather than monitoring changes in cell growth to measure β-lactamase inhibitor efficacy (e.g., traditional methods to determine MICs), our assay provides a more rapid and specific measure of inhibitor potency by measuring the ability of an inhibitor to rescue β-lactam antibiotics from β-lactamase-producing bacteria. This approach can be extended to determine cellular IC_50_ values, allowing for inhibitor potency to be quantified. Our biosensor platform addresses drawbacks associated with assays employing purified β-lactamases that utilize chromogenic (e.g., nitrocefin, CENTA) [10] or fluorogenic (e.g., FC5) [10] substrates. Although assays using purified enzymes are sensitive and rapid, they often require the use of expensive substrates and do not account for cell wall permeability and drug efflux. In Gram-negative bacteria, β-lactamases are located in the periplasm [39], where they degrade β-lactams that have penetrated the outer membrane. Therefore, β-lactamase inhibitors must also reach the periplasm to access their targets. These factors may account for some of the differences between reported IC_50_ values for inhibitors against purified β-lactamases compared to the cellular IC_50_ values determined in this study. For example, several reported NDM-1 inhibitors (DPA, NTA, CAP) have IC_50_ values in the low micromolar range against purified enzymes (Table S3), while these values increased up to > 200-fold when evaluated against cells using the biosensor assay. This highlights the importance of evaluating the efficacy of β-lactamase inhibitors in a cellular context.

Bacterial resistance to β-lactams resulting from the production of β-lactamases continues to present a global health threat due to their ability to hydrolyze all clinically available β-lactams and rapidly overcome β-lactamase inhibitors. The discovery of new inhibitors is imperative in ensuring future clinical utility of β-lactam antibiotics and will be fuelled by the use of assays that prioritize the identification of inhibitors that are effective against bacterial cells. Future work will be aimed at applying this biosensor platform to the screening of small molecule and peptide libraries for the discovery of novel β-lactamase inhibitors and PBP-targeting antibiotics.

## MATERIALS AND METHODS

### Reagents

Imipenem, meropenem, amoxicillin, tazobactam, and clavulanic acid were purchased from Glentham Life Sciences. Avibactam was purchased from Medkoo Biosciences. Dipicolinic acid and nitrilotriacetic acid were purchased from Fisher Scientific. Captopril was purchased from VWR. EDTA was purchased from Thermo Fisher Scientific. Embelin was purchased from Toronto Research Chemicals Inc. 2TY media components (tryptone, yeast extract, sodium chloride) and cation-adjusted Mueller Hinton Broth (CAMHB; Becton Dickinson) were purchased from Fisher Scientific.

### Bacterial culturing

All *Escherichia coli* cultures used for cloning experiments were grown in autoclaved 2TY media (16 g/L tryptone, 10 g/L yeast extract, 5 g/L NaCl) supplemented with the appropriate selection antibiotic, where required. Liquid cultures were incubated at 37 °C and 200 rpm. Overnight plate cultures were grown on 2TY agar plates supplemented with the appropriate selection antibiotic, where required.

### Biosensor development

The *ampR* gene and *ampC* promoter were amplified from the genomic DNA of *P. aeruginosa* 2_1_26 (BEI Resources) by polymerase chain reaction (PCR). The *luxCDABE* cassette was amplified from the pAKgfplux1 plasmid (Addgene # 14083) by PCR. [40] pUC19 vector (New England BioLabs) was digested with SacI (New England BioLabs) and purified by gel extraction following agarose gel electrophoresis. The PCR-amplified *ampR/*P*ampC* and *luxCDABE* sequences were cloned into pUC19 by HiFi assembly (New England BioLabs). Next, the ampicillin resistance marker in this plasmid was disrupted by restriction digest with ScaI (New England Biolabs), and the kanamycin resistance gene from pHSG298 (National BioResource Project, NBRP) was cloned into this site using HiFi assembly to generate the plasmid pAMPLUX. The sequence of pAMPLUX was confirmed by whole plasmid sequencing (Plasmidsaurus). Chemically competent *E. coli* BW25113 (NBRP) cells were transformed with pAMPLUX and subsequently used for biosensor assays.

### Preparation of β-lactamase-producing strains

The TEM-116 gene and promoter were amplified from pUC19 by PCR. Genes and native promoters for NDM-1 (*Klebsiella pneumoniae* AR0041), IMP-1 (*Pseudomonas aeruginosa* AR0103), VIM-2 (*P. aeruginosa* AR0100), KPC-2 (*P. aeruginosa* AR0098), and OXA-48 (*Enterobacter aerogenes* AR0074) were amplified from the genomic DNA of their respective organisms by PCR. All β-lactamase genes (along with their native promoters) were cloned into the HindIII site of pACYC184 (NBRP) by HiFi assembly. The sequences of the recombinant plasmids were confirmed by Sanger sequencing (The Centre for Applied Genomics, The Hospital for Sick Children, Toronto, ON) and whole plasmid sequencing (Plasmidsaurus).

### Minimum inhibitory concentration (MIC) testing

All MIC testing was conducted according to Clinical and Laboratory Standards Institute (CLSI) guidelines [41]. *E. coli* BW25113 cells (untransformed, transformed with pAMPLUX, or transformed with a pACYC184 plasmid encoding a β-lactamase) were cultured overnight at 37 °C on 2TY agar plates supplemented with the appropriate antibiotic, where required. Antibiotic and inhibitor stocks were prepared in sterile water and serially diluted in CAMHB in clear 96-well non-treated flat-bottom microplates (Falcon). Cell suspensions were prepared by resuspending colonies in CAMHB growth media to OD_600_ = 0.1 (approx. 1.5 x 10^8^ CFU/mL). These suspensions were diluted in CAMHB to approx. 5 x 10^6^ CFU/mL, and 20 µL of the suspension was added to 180 µL of each antibiotic/inhibitor treatment, giving a final inoculum of 5 x 10^5^ CFU/mL. Plates were incubated for 16 - 18 hours at 37 °C without shaking, and OD_600_ readings were taken using a Synergy LX plate reader (Agilent BioTek).

### Biosensor bioactivity testing

Biosensor cells (*E. coli* BW25113 transformed with pAMPLUX) were grown overnight at 37 °C on 2TY agar plates supplemented with 50 µg/mL kanamycin. Imipenem and amoxicillin stocks were prepared in sterile water, and 180 µL of each antibiotic was added to a white LUMITRAC 200 96-well plate (Greiner Bio-One) in quadruplicate. Biosensor cell suspensions were prepared to OD_600_ = 0.2 by suspending colonies in 2TY growth media, and 50 µL of suspended cells were added to each imipenem and amoxicillin treatment. Luminescence readings were taken every hour for 3 hours using a Spectramax ID3 plate reader (Molecular Devices); between readings, the plate was incubated at 37 °C.

### Biosensor β-lactamase inhibition assays

*E. coli* BW25113 cells transformed with the pACYC184 plasmids carrying β-lactamase genes were grown overnight at 37 °C on 2TY agar plates supplemented with 25 µg/mL chloramphenicol. Biosensor cells were grown overnight at 37 °C on 2TY agar plates supplemented with 50 µg/mL kanamycin. All β-lactam antibiotics and β-lactamase inhibitors were dissolved in sterile water. Serial dilutions of these solutions were prepared in 2TY, and 180 µL of each treatment was added to white LUMITRAC 200 96-well plates (Greiner Bio-One). Cell suspensions of β-lactamase-producing *E. coli* were prepared by suspending colonies in 2TY growth media to an OD_600_ of 0.2, and 20 µL of each cell suspension was added to each well containing β-lactam antibiotics or β- lactam/β-lactamase inhibitor combinations. Plates were incubated for 1.5 hours at 37 °C without shaking. Biosensor cell suspensions were prepared to OD_600_ = 0.2 by suspending colonies in 2TY growth media, and 50 µL of cells were added to the microplate wells containing β-lactamase-producing cells and β-lactam antibiotics and/or β-lactamase inhibitors. The resulting mixtures were incubated for 2 hours at 37 °C. Luminescence readings were acquired using a Spectramax ID3 multimode plate reader (Molecular Devices). Luminescence values were normalized to the sample which gave the highest luminescence output and expressed as a percentage. The level of β- lactamase activity under the assay conditions was expressed as the % luminescence value subtracted from 100%. IC_50_ values for each inhibitor were determined using the “non-linear curve fit [inhibitor] vs. response” function in GraphPad Prism v9.5.

### Growth-based assays for measuring β-lactamase inhibition

*E. coli* BW25113 cells transformed with pACYC184 plasmids carrying β-lactamase genes were cultured overnight at 37 °C on 2TY agar plates supplemented with 25 µg/mL chloramphenicol. Antibiotic and inhibitor stocks were prepared in sterile water and serially diluted in 2TY growth media in clear 96-well non-treated flat-bottom microplates (Falcon). Cell suspensions were prepared by suspending colonies in 2TY growth media to OD_600_ = 0.2, and 20 µL of this suspension was added to each treatment well containing antibiotics alone or in combination with β-lactamase inhibitors. Plates were incubated for 16 - 18 hours at 37 °C without shaking, and OD_600_ readings were taken using a Synergy LX plate reader (Agilent BioTek).

