## Supplemental Data for "Development of a Whole-Cell Biosensor for β-Lactamase Inhibitor Discovery"

### Supplementary Data: Tables and Figures

**Table S1.** Strains used in this study.

| *E. coli* Strain | Characteristics | Source |
|---|---|---|
| NEB10β | Wild-type | New England BioLabs |
| BW25113 | Wild-type | NBRP |
| BW25113 Δ*ompF* | *ompF* gene disrupted with $kan^R$ marker | Keio Collection[1] (NBRP, Japan) |
| BW25113-Biosensor | Transformed with pAMPLUX | This study |
| BW25113-TEM-116 | Transformed with pACYC184 $bla_{TEM-116}$ | This study |
| BW25113Δ*ompF*-TEM-116 | *ompF* gene disrupted with $kan^R$ marker, transformed with pACYC184 $bla_{TEM-116}$ | This study |
| BW25113-KPC-2 | Transformed with pACYC184 $bla_{KPC2}$ | This study |
| BW25113-OXA-48 | Transformed with pACYC184 $bla_{OXA-48}$ | This study |
| BW25113-NDM-1 | Transformed with pACYC184 $bla_{NDM-1}$ | This study |
| BW25113-IMP-1 | Transformed with pACYC184 $bla_{IMP-1}$ | This study |
| BW25113-VIM-2 | Transformed with pACYC184 $bla_{VIM-2}$ | This study |

**Table S2.** Minimum inhibitory concentration (MIC) values for amoxicillin and meropenem against the strains used in this study.

| ***E. coli*** **Strain** | **MIC (Meropenem, µg/mL)** | **MIC (Amoxicillin, µg/mL)** |
|---|---|---|
| BW25113 | 0.06 | 4 |
| BW25113-Biosensor | 0.06 | 4 |
| BW25113-TEM-116 | 0.06 | > 256 |
| BW25113-KPC-2 | >32 | > 256 |
| BW25113-OXA-48 | 2 | > 256 |
| BW25113-NDM-1 | >32 | > 256 |
| BW25113-IMP-1 | >32 | > 256 |
| BW25113-VIM-2 | 4 | > 256 |

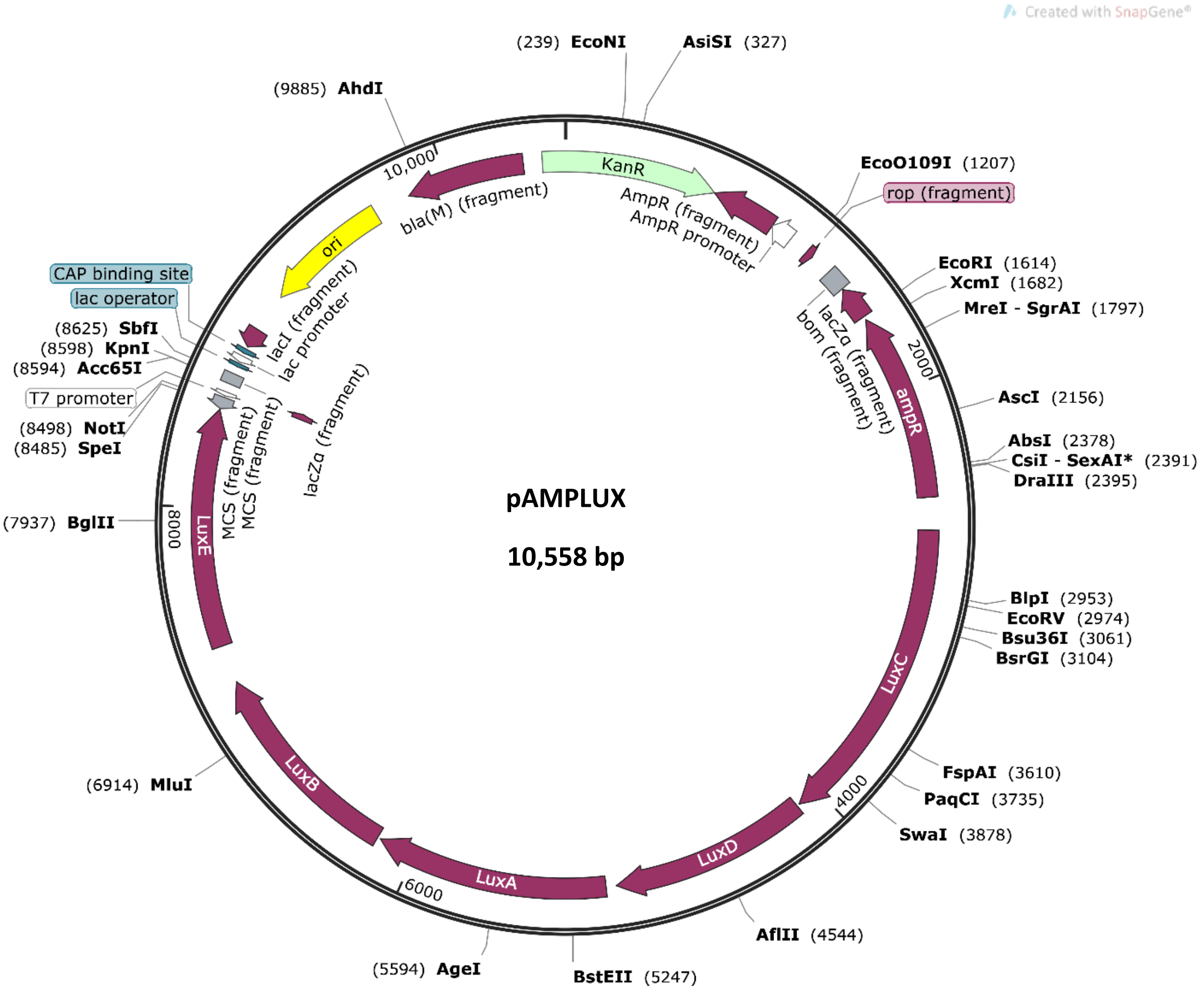

**Figure S1. Plasmid map for pAMPLUX.** The *ampR*-P*ampC* and *luxCDABE* fragments were cloned into pUC19 at the SacI restriction site in the multiple cloning site (MCS). The ampicillin resistance marker (*bla*) in the resulting vector was disrupted by digestion with ScaI, and the kanamycin resistance marker (KanR) was cloned into this site. The plasmid map was prepared by Plasmidsaurus.

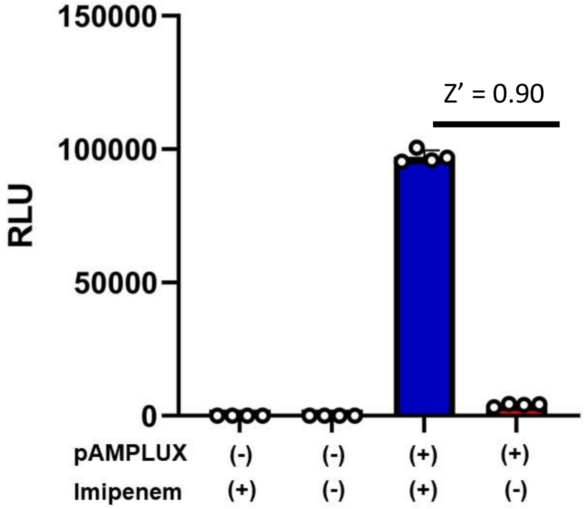

**Figure S2. Verifying specificity of luminescent signal produced by *E. coli* transformed with pAMPLUX in response to β-lactam exposure.** Wild-type *E. coli* BW25113 cells (- pAMPLUX) and cells transformed with the pAMPLUX plasmid (+ pAMPLUX) were incubated in 2TY media supplemented with imipenem (1.6 μM), or 2TY media alone. Luminescence readings were taken after two hours. *E. coli* BW25113 cells that were not transformed with pAMPLUX did not produce luminescence, regardless of whether or not they were exposed to imipenem. *E. coli* cells transformed with pAMPLUX produced luminescence when treated with imipenem, while minimal background luminescence was observed following media treatment alone. To assess the assay performance, the Z-factor (Z') was determined based on the RLU values obtained from the biosensor cells following treatment with imipenem or media alone. Note that a Z' value greater than 0.5 indicates good separation between positive and negative signals.[2]

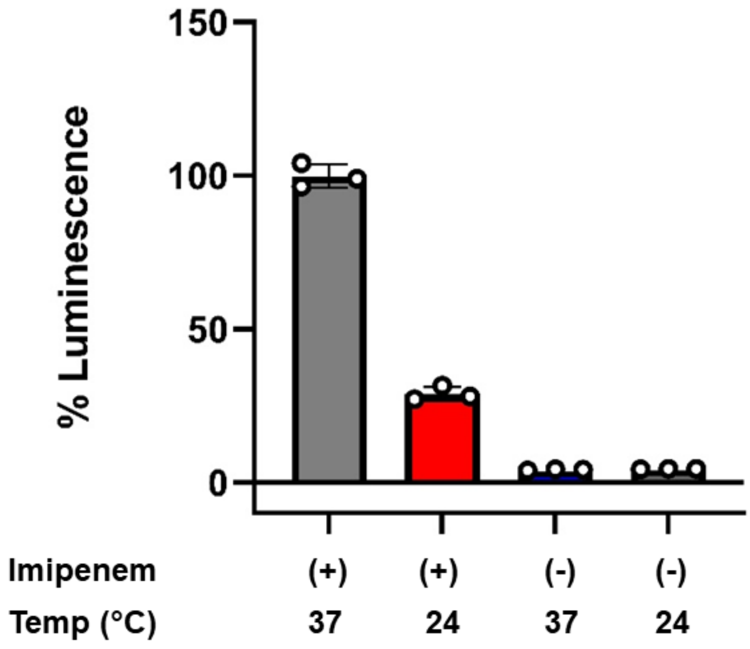

**Figure S3. Performance of the biosensor at different incubation temperatures**. Biosensor cells were treated with either 2TY media supplemented with imipenem (1.6 µM) or 2TY media alone. These cell suspensions were incubated for two hours either at 37 °C or at ambient room temperature (approximately 24 °C), and luminescence was measured. Based on these results, 37 °C was chosen as the temperature for subsequent assays.

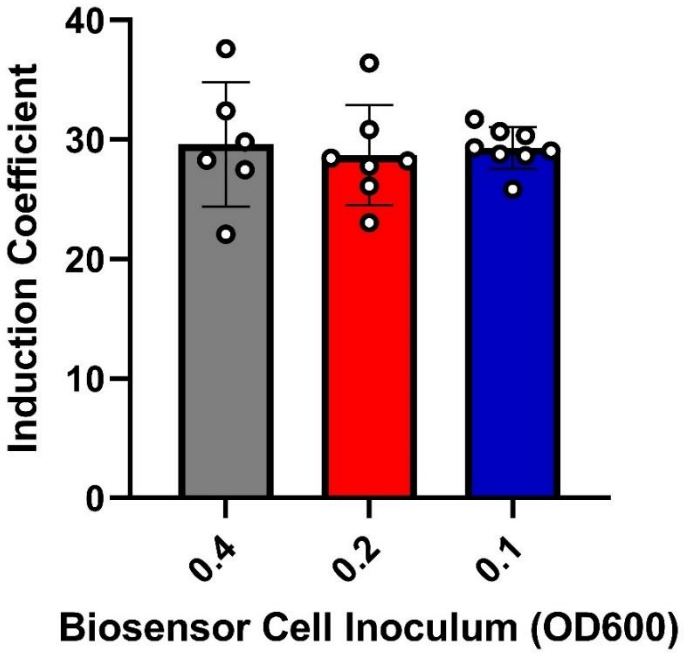

**Figure S4. Impact of biosensor cell density on induction coefficient**. Biosensor cell suspensions were prepared in 2TY media to $OD_{600}$ = 0.4, 0.2, or 0.1, then diluted 10-fold in 2TY supplemented with imipenem (1.6 µM, final concentration). These mixtures were incubated at 37 °C for two hours, and luminescence readings were taken. The induction coefficient was determined by dividing the RLU values obtained from samples treated with imipenem with the RLU values obtained from samples that were treated with 2TY media alone. There was no apparent difference in the induction coefficients determined from the three different biosensor cell suspensions.

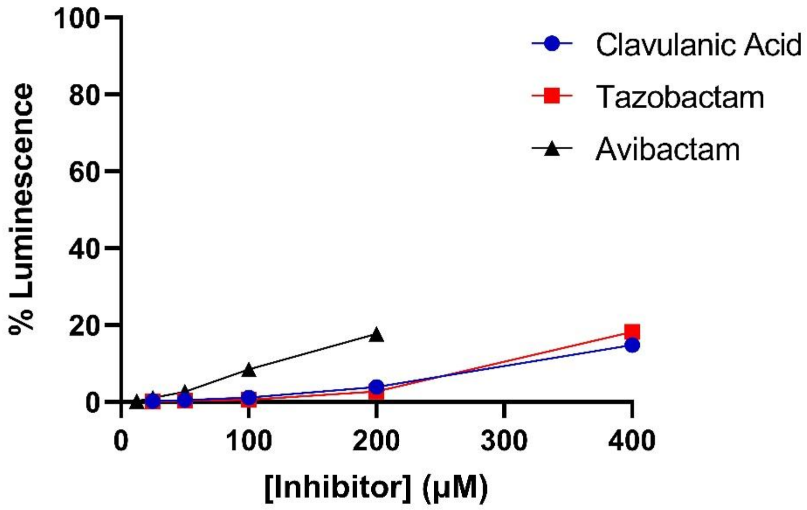

**Figure S5. Induction of biosensor luminescence by SBL inhibitors in the absence of a β-lactam antibiotic.** Biosensor cells were treated with serial dilutions of clavulanic acid, tazobactam or avibactam prepared in 2TY. Mixtures were incubated for two hours, and luminescence readings were taken. The luminescence values were normalized to the luminescence obtained when biosensor cells were treated with amoxicillin (20 μM). n=4, error bars indicate S.D.

**Table S3.** $IC_{50}$ values for inhibitors against β-lactamase-producing cells and against purified β-lactamases.

| Enzyme | Inhibitor | Inhibitor concentration needed to reduce MIC to susceptible range[a,b] | IC50 (μM, Cell)[c] | IC50 (μM, Enzyme)[d] |
|---|---|---|---|---|
| TEM-116 | Tazobactam[e] | 200 | 118 ± 32.2 | 0.13[3] [j] |
| | Clavulanic acid[e] | 100 | 64.2 ± 18.7 | 0.08[3] |
| | Avibactam[e] | 30 | 13.2 ± 2.1 | 0.008[4] |
| KPC-2 | Tazobactam[e] | N.I.[m] | N.I.[m] | N.I.[m] |
| | Clavulanic acid[e] | N.I.[m] | N.I.[m] | N.I.[m] |
| | Avibactam[e] | 60 | 29.2 ± 10.1 | 0.17[4] |
| OXA-48 | Tazobactam[e] | N.I.[m] | N.I.[m] | N.I.[m] |
| | Clavulanic acid[e] | N.I.[m] | N.I.[m] | N.I.[m] |
| | Avibactam[e] | 60 | 35.5 ± 4.6 | 0.18[5] |
| NDM-1 | Dipicolinic acid[f] | 300 | 238.5 ± 4.5 | 3.8[6] |
| | Nitrilotriacetic acid[f] | 600 | 383.7 ± 6.3 | 1.3[6] |
| | Ethylenediaminetetraacetic acid[f] | 40 | 11.4 ± 0.6 | 0.052[7] |
| | Captopril[f] | 600 | 461 ± 47 | 20.1[8] |
| | Embelin[f] | 300 | 146 ± 13 | 2.1[9] |
| IMP-1 | Dipicolinic acid[f] | 300 | 243.8 ± 4.2 | N.R.[n] |
| | Nitrilotriacetic acid[f] | N.I.[m] | N.I.[m] | N.R.[n] |
| | Ethylenediaminetetraacetic acid[f] | 200 | 51.8 ± 5.6 | 55[10] |
| | Captopril[f] | 600 | 299 ± 19 | 7.1[8] |
| | Embelin[f] | N.I.[m] | N.I.[m] | 100[9] |
| VIM-2 | Dipicolinic acid[g] | N.T.[h,o] | 210.3 ± 10.2 | 2.9[6] |
| | Nitrilotriacetic acid[g] | N.T.[h,o] | 331 ± 46 | 2.4[6] |
| | Ethylenediaminetetraacetic acid[g] | N.T.[h,o] | 68.2 ± 22.5 | 9.3[k]-200[11, 12] |
| | Captopril[g] | N.T.[h,o] | 145 [i] | 0.07[8] |
| | Embelin[g] | N.I.[h] | N.I.[m] | 200[9] [l] |

[a] The susceptibility breakpoints for meropenem and amoxicillin are 1 μg/mL and 4 μg/mL, respectively; [b] measured based on cell growth after 18 hours using the same conditions as biosensor assays;[c] Cell-based $IC_{50}$ values determined in this study using the biosensor. All inhibitors were tested against *E. coli* BW25113 cells producing the β-lactamase indicated; [d] Inhibitor $IC_{50}$ values reported in literature, references are indicated with superscript numbers; [e] administered with 20 μM amoxicillin; [f] administered with 2.5 μM meropenem; [g] administered with 650 nM meropenem; [h] did not test with breakpoint concentration of meropenem due to poor VIM-2 expression in *E. coli*, resulting from inefficient periplasmic processing of

the enzyme[13]; [i] could not be accurately determined due to the shape of the dose response curve; [j] reported $IC_{50}$s are for TEM-1, not TEM-116; [k] $IC_{50}$ for EDTA reported for VIM-1; [l] IC50 for embelin was only reported for VIM-1, not VIM-2. [m]N.I. = not inhibited; [n]N.R. = not reported; [o]N.T. = not tested.

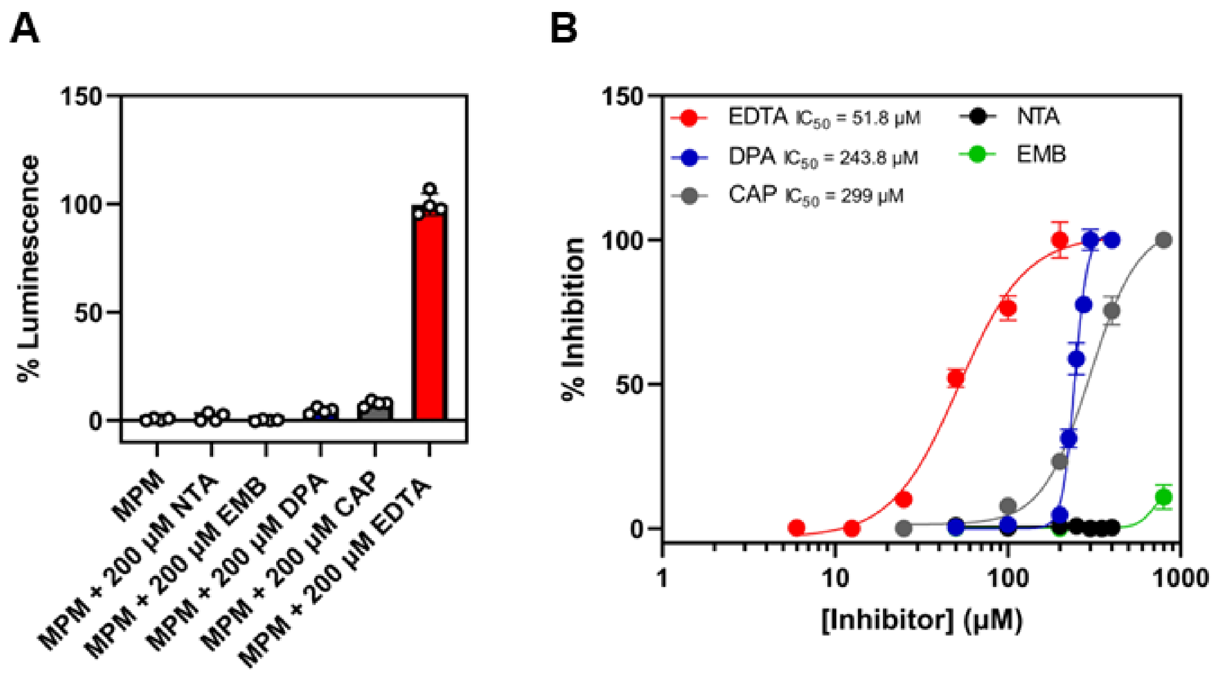

**Figure S6. IMP-1 inhibition assays.** (A) Single point and (B) dose response analysis of IMP-1 inhibition. IMP-1-producing *E. coli* were treated with meropenem (MPM; 2.5 µM) in combination with ethylenediaminetetraacetic acid (EDTA), dipicolinic acid (DPA), nitrilotriacetic acid (NTA), captopril (CAP) or embelin (EMB) at the concentrations indicated. Luminescence readings were normalized to the sample with the greatest luminescence to determine % inhibition. n=4, error bars indicate S.D.

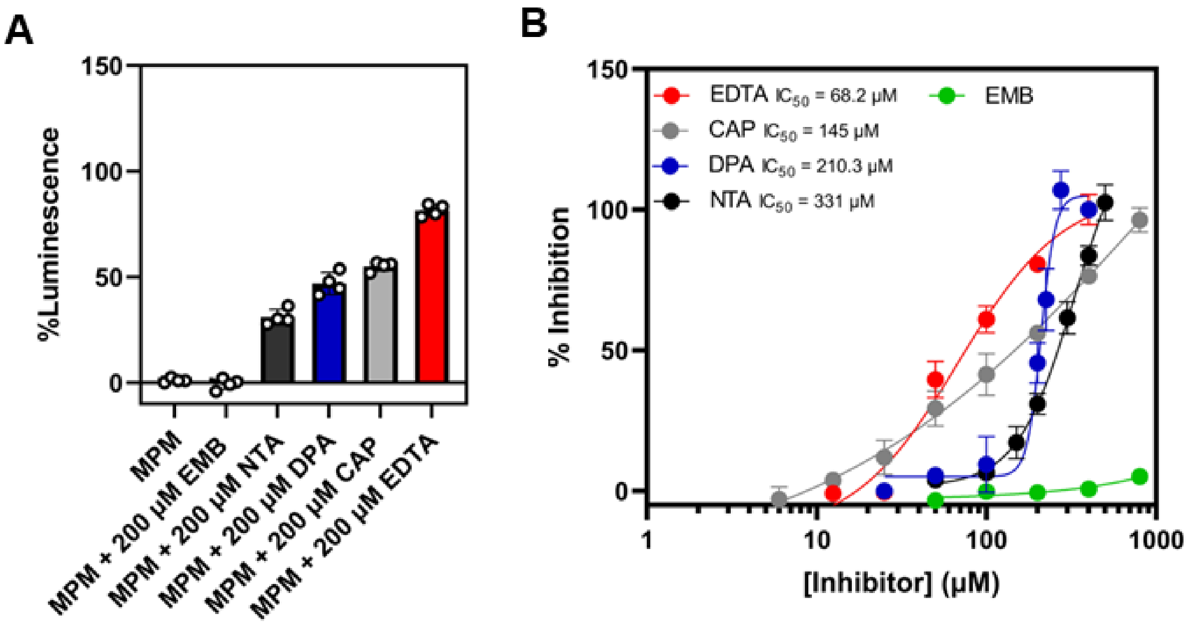

**Figure S7. VIM-2 inhibition assays.** (A) Single point and (B) dose response analysis of VIM-2 inhibition. VIM-2-producing *E. coli* were treated with meropenem (MPM; 650 nM) in combination with ethylenediaminetetraacetic acid (EDTA), dipicolinic acid (DPA), nitrilotriacetic acid (NTA), captopril (CAP) or embelin (EMB) at the concentrations indicated. Luminescence readings were normalized to the sample with the greatest luminescence to determine % inhibition. n=4, error bars indicate S.D.

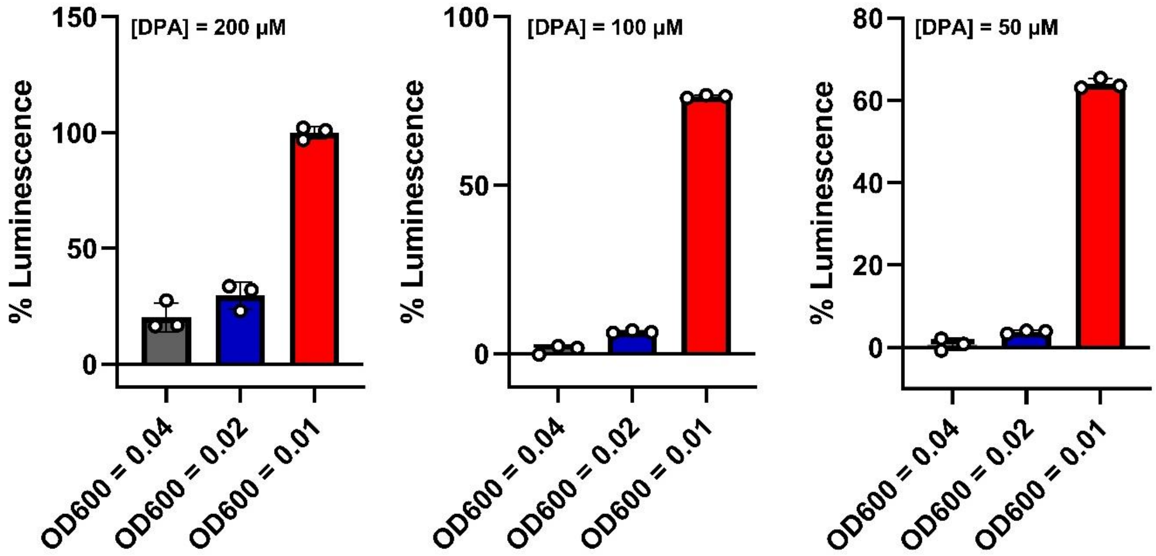

**Figure S8. Effects of β-lactamase culture inoculum on apparent potency of dipicolinic acid (DPA) against NDM-1 in *E. coli* cells**. Suspensions of NDM-1-producing *E. coli* were prepared to OD600 = 0.4, 0.2 or 0.1, then diluted ten-fold in 2TY supplemented with meropenem (final concentration 2.5 µM) and DPA. Luminescence readings were normalized to the sample with the greatest luminescence. n=4, error bars indicate S.D.

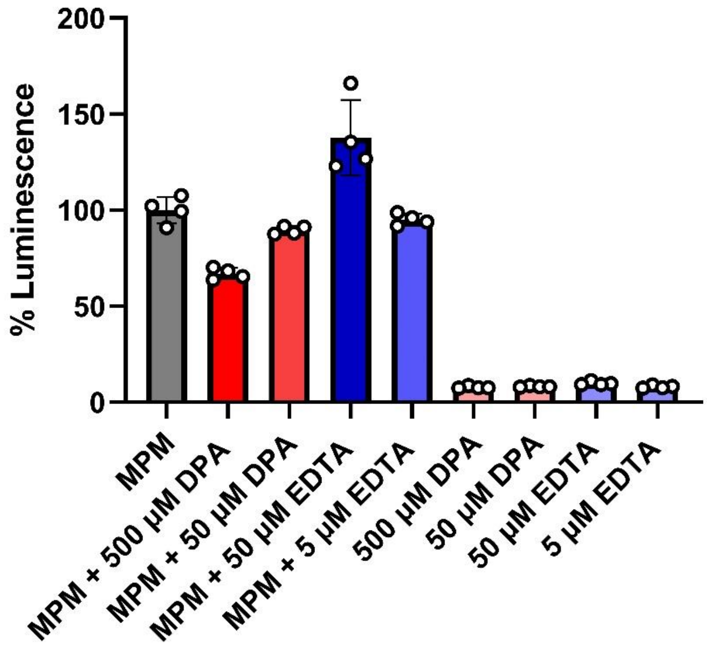

**Figure S9. Impact of EDTA and DPA on biosensor induction.** Biosensor cells were treated with meropenem (MPM; 2.5 µM) alone or in combination with several concentrations of ethylenediaminetetraacetic acid (EDTA) or dipicolinic acid (DPA). Elevated luminescence readings for MPM + 50 µM EDTA indicate increased antibiotic entry due to permeabilization of the outer membrane by EDTA. Lower luminescence readings observed for MPM + 500 µM DPA is indicative of biosensor cell death due to permeabilization of the outer membrane by DPA. Luminescence readings were normalized to the meropenem only treatment. n=4, error bars indicate S.D.
